# Leveraging single-cell ATAC-seq to identify disease-critical fetal and adult brain cell types

**DOI:** 10.1101/2021.05.20.445067

**Authors:** Samuel S. Kim, Karthik Jagadeesh, Kushal K. Dey, Amber Z. Shen, Soumya Raychaudhuri, Manolis Kellis, Alkes L. Price

**Affiliations:** Department of Electrical Engineering and Computer Science, Massachusetts Institute of Technology, Cambridge, MA, 02142; Department of Epidemiology, Harvard T.H. Chan School of Public Health, Boston, MA, 02115; Department of Mathematics, Massachusetts Institute of Technology, Cambridge, MA, 02142; Division of Genetics, Department of Medicine, Brigham and Women’s Hospital and Harvard Medical School, Boston, MA 02115; Program in Medical and Population Genetics, Broad Institute of MIT and Harvard, Cambridge, MA, 02142; Department of Biostatistics, Harvard T.H. Chan School of Public Health, Boston, MA, 02115

## Abstract

Prioritizing disease-critical cell types by integrating genome-wide association studies (GWAS) with functional data is a fundamental goal. Single-cell chromatin accessibility (scATAC-seq) and gene expression (scRNA-seq) have characterized cell types at high resolution, and early work on integrating GWAS with scRNA-seq has shown promise, but work on integrating GWAS with scATAC-seq has been limited. Here, we identify disease-critical fetal and adult brain cell types by integrating GWAS summary statistics from 28 brain-related diseases and traits (average *N* =298K) with 3.2 million scATAC-seq and scRNA-seq profiles from 83 cell types. We identified disease-critical fetal (resp. adult) brain cell types for 22 (resp. 23) of 28 traits using scATAC-seq data, and for 8 (resp. 17) of 28 traits using scRNA-seq data. Notable findings using scATAC-seq data included highly significant enrichments of fetal photoreceptor cells for major depressive disorder, fetal ganglion cells for BMI, fetal astrocytes for ADHD, and adult VGLUT2 excitatory neurons for schizophrenia. Our findings improve our understanding of brain-related diseases and traits, and inform future analyses of other diseases/traits.

## Introduction

Genome-wide association studies (GWAS) have been successful in identifying disease-associated loci, occasionally producing valuable functional insights ^1,2^. Identifying disease-critical cell types is a fundamental goal for understanding disease mechanisms, designing functional follow-ups, and developing disease therapeutics^3^. Several studies have identified disease-critical tissues and cell types using bulk chromatin ^4–9^ and/or gene expression data ^8,10–12^.

With the emergence of single-cell profiling of diverse tissues and cell types ^13–17^, several studies have integrated GWAS data with single-cell chromatin accessibility (scATAC-seq) ^16–19^ and single-cell gene expression (scRNA-seq) ^10,20,21^. However, compared to scRNA-seq data, scATAC-seq data has been less well-studied for identifying disease-critical cell types. In addition, while it is widely known that biological processes in the human brain vary with developmental stage ^22–26^, the impact on disease risk of cell types in different developmental stages of the brain has not been widely explored.

Here, we sought to infer disease-critical cell types by analyzing scATAC-seq and scRNA-seq data derived from single-cell profiling of over 3 million cells from fetal and adult human brains. We analyzed 83 brain cell types from 4 single-cell datasets ^14–17^ across 28 brain-related diseases and complex traits (average *N* = 298K). We determined that both scATAC-seq and scRNA-seq data were highly informative for identifying disease-critical cell types; surprisingly, scATAC-seq data was somewhat more informative in the data that we analyzed.

## Methods

### 28 distinct brain-related diseases and traits

We considered 146 sets of GWAS summary association statistics, including 83 traits from the UK Biobank and 63 traits from publicly available sources, with z-scores for total SNP-heritability of at least 6 (computed using S-LDSC with the baseline-LD (v.2.2) model). We selected 31 brain-related traits based on previous studies^8,17,20,21,27^. We removed 3 traits (with lower SNP-heritability z-score) that had a genetic correlation of at least 0.9 with at least one of these 31 traits, retaining a final set of 28 distinct brain-related traits (including 7 traits from the UK Biobank) (Table S1). The genetic correlations among the 28 traits are reported in Table S2.

We additionally analyzed 6 distinct control (non-brain-related) traits: coronary artery disease, bone mineral density, rheumatoid arthritis, type 2 diabetes, sunburn occasion, and breast cancer. These 6 traits had similar sample sizes and SNP-heritability z-scores as the 28 brain-related traits.

### Genomic annotations and the baseline model

We define a binary genomic annotation as a subset of SNPs in a predefined reference panel. We restrict our analysis to SNPs with a minor allele frequency (MAF) ≥ 0.5% in 1000 Genomes ^28^ (see Web resources).

The baseline model ^29^ (v.1.2; see Table S3) contains 53 binary functional annotations (see Web resources). These annotations include genomic elements (e.g. coding, enhancer, UTR), regulatory elements (e.g. histone marks), and evolutionary constraint. We included the baseline model, consistent with ref. ^8,30^, when assessing the heritability enrichment of the cell-type annotations.

### Single-cell ATAC-seq data

We considered single-cell ATAC-seq data for fetal brains from Domcke et al. ^16^ (donor size = 26) and adult brains (isocortex, striatum, hippocampus, and substantia nigra) of cognitively healthy individuals from Corces et al. ^17^ (donor size = 10). We used the chromatin accessible peaks for each cell type without modifications (see Web resources). In short, these peaks refer to MACS2^31^ peak regions, excluding the ENCODE blacklist regions. For the Domcke et al. data, authors called peaks on each tissue sample and then generated a masterlist of all peaks across all samples and generated the cell-type-specific peaks using Jensen-Shannon divergence^32^. To further ensure the cell-type specificity, we used the union of per-dataset open chromatin regions across all cell types as the background annotation in the S-LDSC conditional analysis.

### Single-cell RNA-seq data analyzed

We considered single-cell RNA-seq data for fetal brains from Cao et al. ^15^ (donor size = 28) and single-cell RNA-seq data for non-fetal brains (prefrontal cortex and anterior cingulate cortex) from Velmeshev et al. ^14^ (donor size = 31). For Cao et al. data, we processed data from three brain-related organs: cerebellum, cerebrum, and eye. We followed the processing protocol as outlined in ref. 21. For each data set, we downloaded metadata for each cell including the total number of reads and sample ID. We then transformed each expression matrix to log2(TP10K+1) unit. We performed a dimensional reduction using a principal component analysis with the top 2,000 highly variable genes, batch correction using Harmony ^33^, and applied the Leiden graph clustering method ^34^.

To obtain specifically expressed (SE) gene scores for each cell type, we applied a non-parametric Wilcoxon rank-sum test between gene expression from focal cell type vs. gene expression in other cell types, as in ref. 21; specific expression was assessed relative to all brain cell types. We transformed the per-gene SE p-value to lie between 0 and 1, by applying min-max normalization on -2log(p-value), indicating a relative importance of each gene in each cellular process.

To construct SNP annotations from gene scores, we employed an enhancer-gene linking strategy from the union of the Roadmap^7^ and Activity-By-Contact (ABC^35,36^) strategies, as in ref. 21. Because we focused on brain-related traits, we used brain-specific enhancer-gene links. SNP annotation values were defined based on the maximum gene score among genes linked to a SNP (or 0 when no genes are linked to a SNP).

### Enrichment and *τ* * metrics

We used stratified LD score regression (S-LDSC^11,29^) to assess the contribution of an annotation to disease and complex trait heritability.

Let *a*_*cj*_ represent the (binary or probabilistic) annotation value of the SNP *j* for the annotation *c*. S-LDSC assumes the variance of per normalized genotype effect sizes is a linear additive contribution to the annotation *c*:

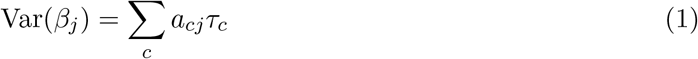

where *τ*_*c*_ is the per-SNP contribution of the annotation *c*. S-LDSC estimates *τ*_*c*_ using the following equation:

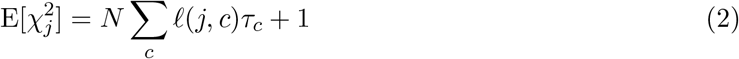

where *N* is the sample size of the GWAS and 𝓁(*j, c*) is the LD score of the SNP *j* to the annotation *c*. The LD score is computed as follow 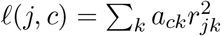 where *r*_*jk*_ is the correlation between the SNPs *j* and *k*.

We used two metrics to assess the informativeness of an annotation. First, the standardized effect size (*τ*\*), the proportionate change in per-SNP heritability associated with a one standard deviation increase in the value of the annotation (conditional on all the other annotations in the model), is defined as follows:

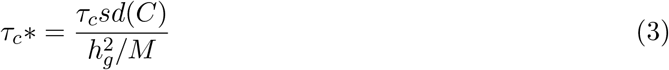

where *sd*(*C*) is the standard deviation of the annotation *c*, 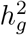 is the estimated SNP-heritability, and *M* is the number of variants used to compute 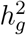 (in our experiment, *M* is equal to 5,961,159, the number of common SNPs in the reference panel). The significance for the effect size for each annotation, as mentioned in previous studies ^27,29,37^, is computed as 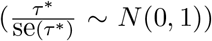, assuming that 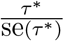 follows a normal distribution with zero mean and unit variance.

Second, enrichment of the binary and probabilistic annotation is the fraction of heritability explained by SNPs in the annotation divided by the proportion of SNPs in the annotation, as shown below:

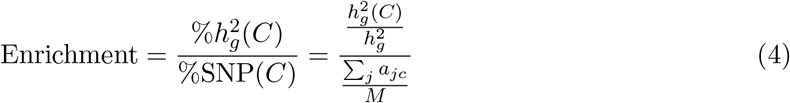

where 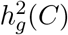 is the heritability captured by the *c*th annotation. When the annotation is enriched for trait heritability, the enrichment is *>* 1; the overlap is greater than one would expect given the trait heritablity and the size of the annotation. The significance for enrichment is computed using the block jackknife as mentioned in previous studies^8,11,27,37^.). The key difference between enrichment and *τ* * is that *τ* * quantifies effects that are unique to the focal annotation after conditioning on all the other annotations in the model, while enrichment quantifies effects that are unique and/or non-unique to the focal annotation.

We used European samples in 1000G ^28^ as reference SNPs and HapMap 3^38^ SNPs as regression SNPs (see Web resources). We excluded SNPs with marginal association statistics *>* 80 and SNPs in the major histocompatibility complex region. In all our analyses, we used the p-value of *τ* * as our primary metric to estimate the effect sizes conditional on known annotations (by including the baseline model as recommended previously^8,30^). We excluded trait-annotation pairs with negative *τ* *, consistent with previous studies ^16,27,39^. We assessed the statistical significance of trait-cell type associations based on per-dataset FDR *<* 5%, aggregating across 28 brain-related traits and all cell types in the dataset (or aggregating across 6 control traits and all cell types in the dataset, in analyses of control traits). As we expect no enrichments of brain cell types in these 6 control traits, we controlled FDR separately from the analysis of brain traits.

### Gene set enrichment analysis using GREAT

We performed gene set enrichments on each cell-type annotations for the gene ontology (GO) biological process, cellular component, and molecular function. We used GREAT^40^ (v.4.0.4) with its default setting, where each gene is assigned a regulatory domain (for proximal: 5kb upstream, 1kb downstream of the TSS; for distal: up to 1Mb). Because annotations from the scRNA-seq were probabilistic, we limited to regions with gene membership probability *>*= 0.8 for gene set enrichment analysis. We used all regions for the scATAC-seq annotations as an input. We defined significant results as those with the FDR-corrected binomial test p-value *<* 0.05.

## Results

### Overview of methods

We define a *cell-type annotation* as an assignment of a binary or probabilistic value between 0 and 1 to each SNP in the 1000 Genomes European reference panel^28^, representing the estimated contribution of that SNP to gene regulation in a particular cell type. Here, we constructed cell-type annotations for 4 datasets: (1) fetal brain scATAC-seq ^16^ (number of cell types (*C*) = 14), (2) fetal brain scRNA-seq data^15^ (*C* = 34), (3) adult brain scATAC-seq ^17^ (*C* = 18), and (4) adult brain scRNA-seq data ^14^ (*C* = 17) (see Web resources).

For scATAC-seq cell-type annotations, we used the chromatin accessible peaks (MACS2^31^ peak regions) provided by ref. ^16,17^. These peaks correspond to accessible regions for transcription factor binding, indicative of active gene regulation. For scRNA-seq cell-type annotations, we annotated SNPs linked to specifically expressed genes in a given cell type ^8^ (compared to other brain cell types) using brain-specific enhancer-gene links ^7,21,35,36^.

We assessed the heritability enrichments of the resulting cell-type annotations by applying S-LDSC ^11^ across 28 distinct brain-related diseases and traits (pairwise genetic correlation *<* 0.9; average *N* =298K; Table S1) to identify significant disease-cell type associations (Figure S1). For each disease-cell type pair, we estimated the heritability enrichment^11^ (the proportion of heritability explained divided by the proportion of annotated SNPs) and standardized effect size ^29^ (*τ* *, defined as the proportionate change in per-SNP heritability associated to a one standard deviation increase in the value of the annotation, conditional on other annotation). We assessed the statistical significance of disease-cell type associations based on per-dataset FDR *<* 5% (for each of 4 datasets, aggregating diseases and cell types) based on p-values for positive *τ* *, as *τ* * quantifies effects that are unique to the cell-type annotation. We conditioned the analyses on a broad set of coding, conserved, and regulatory annotations from the baseline model ^11^ (Table S3). For scATAC-seq annotations, we additionally conditioned on the union of open chromatin regions across all brain cell types in each data set analyzed (consistent with ref. 39, 41), a conservative step to ensure cell-type specificity (see Discussion). For scRNA-seq annotations, we additionally conditioned on the union of brain-specific enhancer-gene links across all genes analyzed (consistent with ref. 21). We did not condition on the LD-related annotations included in the baseline-LD model of ref. ^29,42^, as these annotations reflect the action of negative selection, which may obscure cell-type-specific signals ^30^. Further details are provided in the Methods section. We have publicly released all cell-type annotations analyzed in this study and source code for all primary analyses (see Data and code availability).

### Identifying disease-critical cell types using fetal brain data

We sought to identify disease-critical cell types using fetal brain data, across 28 distinct brain-related diseases and traits (Table S1). We analyzed 14 fetal brain cell types from scATAC-seq data ^16^ (donor size = 26; fetal age of 72-129 days) and 34 fetal brain cell types from scRNA-seq data ^15^ (donor size = 28; fetal age of 89-125 days) (Table S4; see Methods).

We first analyzed fetal brain scATAC-seq data spanning 14 cell types ^16^. We identified 152 significant disease-cell type pairs (FDR *<* 5% for positive *τ* * conditional on other annotations; Table 1, Table 2, Figure 1A, Table S5). Consistent with previous genetic studies ^8,17,20^, we identified strong enrichments of excitatory (i.e. glutamatergic) neurons in psychiatric and neurological disorders, including schizophrenia (SCZ), major depressive disorder (MDD), and attention deficit hyperactivity disorder (ADHD) (Figure 1A). In particular, the role of glutamatergic neurons in MDD is well-supported, as evident from decreased glutamatergic neurometabolite levels in subjects with depression ^43^.

**Table 1.**
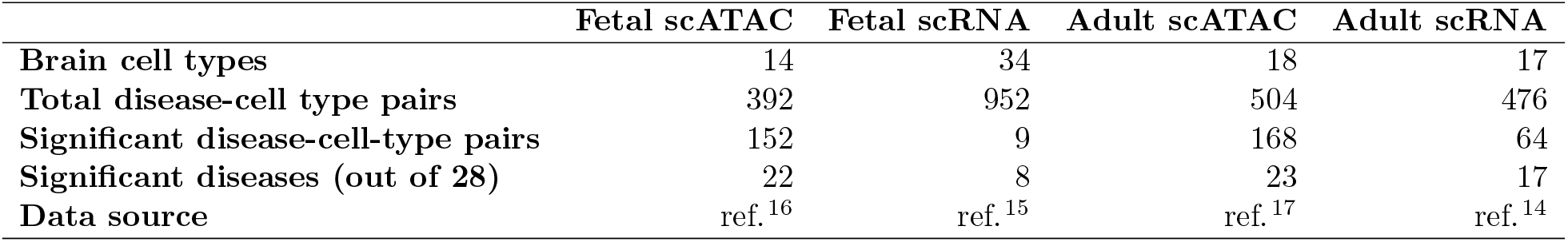
Summary of findings. For each of 4 single-cell chromatin and gene expression data sets analyzed, we report the number of brain cell types analyzed, the total number of disease-cell type pairs analyzed (based on 28 diseases/traits), the number of significant disease-cell type pairs (FDR *<* 5% for positive *τ* *), and the number of diseases/traits with a significant disease-cell type pair.

**Table 2.**
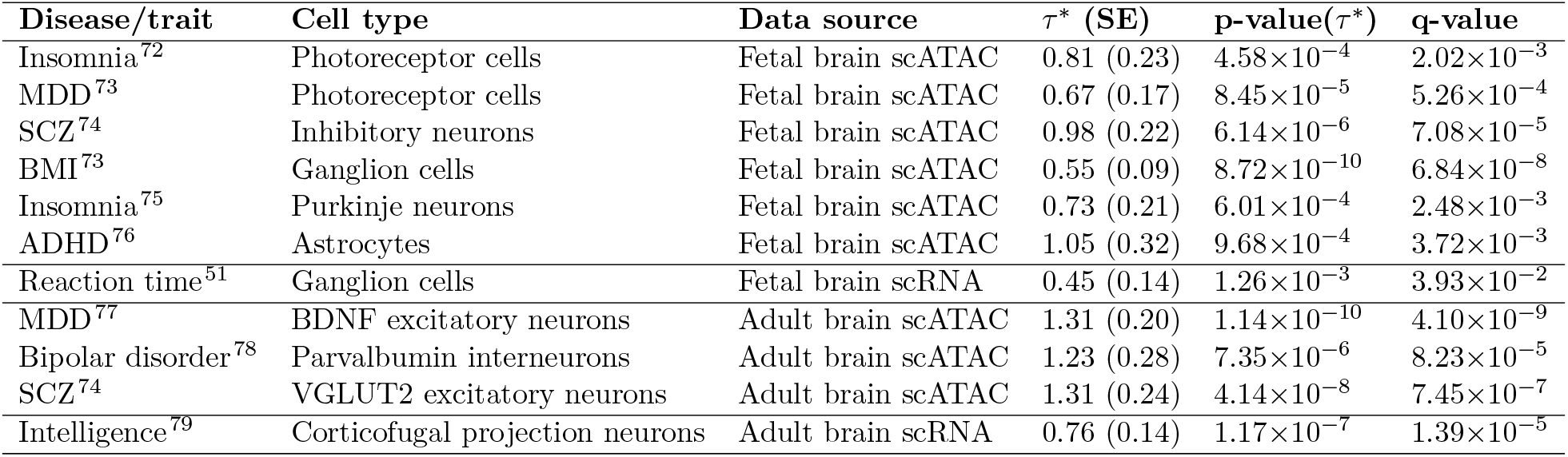
Notable disease-cell type associations. We report the disease/trait, cell type, data source, standardized effect size (*τ* *), p-value for positive *τ* *, and FDR q-value for selected results. Full results are reported in Table S5, Table S7, Table S15, Table S16. A description of diseases/traits analyzed is provided in Table S1. MDD: major depressive disorder, SCZ: schizophrenia, BMI: body mass index, ADHD: attention deficit hyperactivity disorder.

**Figure 1.**
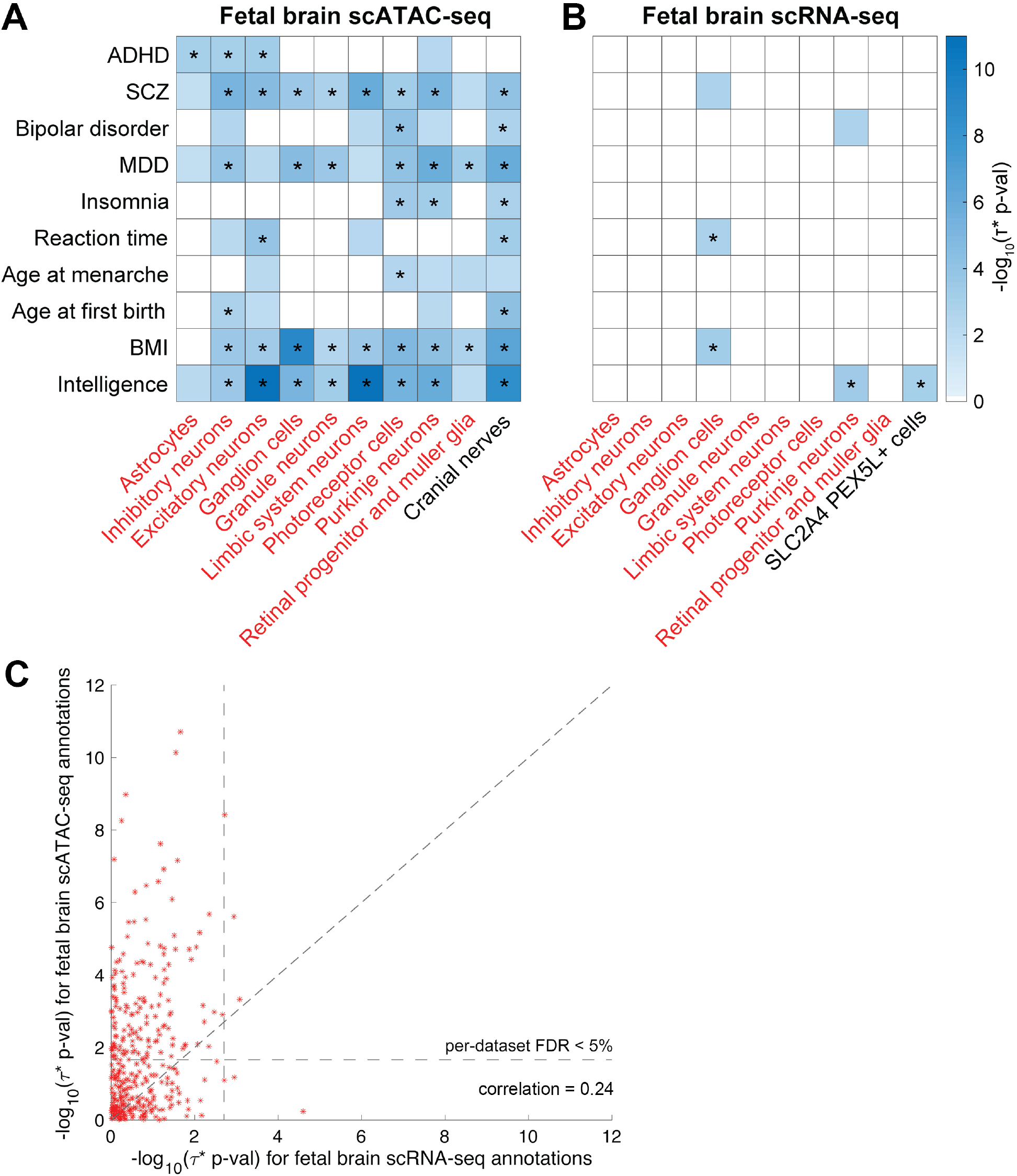
Disease enrichments of cell-type annotations derived from fetal brain. We report (A) −log_10_ p-values for positive *τ* * for a subset of 10 (of 28) diseases/traits and 10 (of 14) fetal brain scATAC-seq cell type annotations; (B) −log_10_ p-values for positive *τ* * for a subset of 10 (of 28) diseases/traits and 10 (of 34) fetal brain scRNA-seq cell type annotations; and (C) comparison of results for 13 cell types included in both fetal brain scATAC-seq and scRNA-seq data. In A-B, only statistically significant results (FDR *>* 5%) are colored (−log_10_(p-value) ≥ 1.67 for scATAC-seq, ≥ 2.70 for scRNA-seq). In A-B, cell types appearing in both datasets are denoted in red font. Numerical results for all diseases/traits and cell types are reported in Table S5, Table S7, and Table S8. * denotes Bonferroni-significant results. ADHD: attention deficit hyperactivity disorder, SCZ: schizophrenia, MDD: major depressive disorder, BMI: body mass index.

Our results also highlight several disease-cell type associations that have not (to our knowledge) previously been reported in analyses of genetic data (Table 2). First, photoreceptor cells were enriched in insomnia. Photoreceptor cells, present in the retina, convert light into signals to the brain, and thus play an essential role in circadian rhythms ^44^, explaining their potential role in insomnia. Second, photoreceptor cells were also enriched in MDD, a genetically uncorrelated trait (*r* = -0.01 with insomnia) (as well as neuroticism; *r* = 0.68 with MDD). Recent studies support the relationship between the degeneration of photoreceptors and anxiety and depression ^45^, suggesting that mental illness is not only a consequence but also a cause of visual impairment. Third, inhibitory (GABAergic) neurons were enriched in SCZ, consistent with GABA dysfuction in the cortex of schizophrenics ^46^. Fourth, ganglion cells were enriched in BMI. Ganglion cells are the projection neurons of the retina, relaying information from bipolar and amacrine cells to the brain. Patients with morbid obesity display significant differences in retinal ganglion cells, retinal nerve fiber layer thickness, and choroidal thickness^47^. Fifth, purkinje neurons were enriched in insomnia (as well as sleep duration (*r* = -0.03 with insomnia) and chronotype (*r* = -0.03 with insomnia; *r* = -0.01 with sleep duration)). While purkinje neurons play a major role in controlling motor movement, they also regulate the rhythmicity of neurons, consistent with a role in impacting sleep^48^. Sixth, astrocytes were enriched in ADHD. Astrocytes perform various functions including synaptic support, control of blood flow, and axon guidance ^49^. In particular, ref. ^50^ highlighted the role of the astrocyte Gi-coupled GABA_B_ pathway activation resulting in ADHD-like behaviors in mice.

We next analyzed fetal brain scRNA-seq data spanning 34 cell types ^15^ (of which 13 were also included in fetal brain scATAC-seq data; Table S6). We identified 9 significant disease-cell type pairs (FDR *<* 5% for positive *τ* * conditional on other annotations; Table 1, Table 2, Figure 1B, Table S7). When restricting to the 7 significant disease-cell type pairs corresponding to the 13 cell types included in both scATAC-seq and scRNA-seq data, 6 of 7 were also significant in analyses of scATAC-seq data. In particular, the enrichment of retinal ganglion cells in reaction time (*p* = 1.26 ×10^−3^ in scRNA-seq data, FDR *q* = 0.039) was non-significant in scATAC-seq data (*p* = 0.028, FDR *q* = 0.060). The enrichment of retinal ganglion cells in reaction time has not (to our knowledge) previously been reported in analyses of genetic data. Previous genetic analyses have focused on enrichments of cerebellum and brain cortex in reaction time ^51^, but the involvement of retinal ganglion cells in receiving visual information and propagating it to the rest of the brain is consistent with a role in visual reaction time^52^.

We compared the results for 13 fetal brain cell types included in both the scATAC-seq and scRNA-seq datasets (Figure 1C and Table S8). While scATAC-seq and scRNA-seq cell-type annotations for matched cell types were approximately uncorrelated to each other (*r* = 0.01 -0.06; Table S9), the corresponding −log_10_(p-values) for positive *τ* * were moderately correlated (*r* = 0.24), confirming the shared biological information. We observed more significant p-values for scATAC-seq than for scRNA-seq in these data sets (see Discussion).

We performed 4 secondary analyses. First, we analyzed enrichments of both scATAC-seq and scRNA-seq brain cell types in 6 control (non-brain-related) diseases and complex traits. As expected, we did not identify any significant enrichments (Table S10 and Table S11). Second, we performed gene set enrichment analysis using GREAT^40^ for both scATAC-seq and scRNA-seq cell-type annotations. As expected, we identified significant enrichments in relevant gene sets (e.g.”photoreceptor cell differentiation” for photoreceptor cells from scATAC-seq; “negative regulation of cell projection organization” for ganglion cells from scRNA-seq; Table S12). Third, for the fetal scRNA-seq data^15^, we constructed annotations based on a +/-100kb window-based strategy (previously used in ref. ^8^) instead of brain-specific enhancer-gene links^7,35,36^ (used in ref. ^21^). We identified 22 significant disease-cell type pairs (Table S13), vs. only 9 using brain-specific enhancer-gene links (although we observed a much stronger opposite trend in adult scRNA-seq data; see below). Fourth, we analyzed bulk chromatin data (7 chromatin marks) spanning 5 fetal brain tissues^9^ (age 52-142 days). We identified 541 significant disease-tissue-chromatin mark triplets spanning 26 of 28 brain-related traits (Table S14). These results are included for completeness, but cannot achieve the same cell-type specificity as analyses of single-cell data.

### Identifying disease-critical cell types using adult brain data

We sought to identify disease-critical cell types using adult brain data, across 28 distinct brain-related diseases and traits (Table S1). Analysis of brains with varying developmental stages might elucidate biological mechanisms, as brains undergo changes in cell type composition and gene expression during development^26,53^. We analyzed 18 adult brain cell types from scATAC-seq data ^17^ (donor size = 10; age 38-95 years) and 17 adult brain cell types from scRNA-seq data ^14^ (donor size = 31; age 4-22 years) (Table S4; see Methods). For brevity, we use the term “adult” to refer to child and adult donors who have surpassed the fetal development stage.

We first analyzed adult brain scATAC-seq data spanning 18 cell types ^17^. We identified 168 significant disease-cell type pairs (FDR *<* 5% for positive *τ* * conditional on other annotations; Table 1, Table 2, Figure 2A, Table S15). Consistent with previous genetic studies ^8,17,19,39^, we identified strong enrichments of excitatory neurons in SCZ and bipolar disorder (genetic correlation = 0.70) (Figure 2A). Although an analysis of mouse scATAC-seq identified a significant enrichment of excitatory neurons in SCZ cases vs. bipolar cases ^19^, we did not replicate this finding (*p* = 0.66 for positive *τ* *; Table S15).

**Figure 2.**
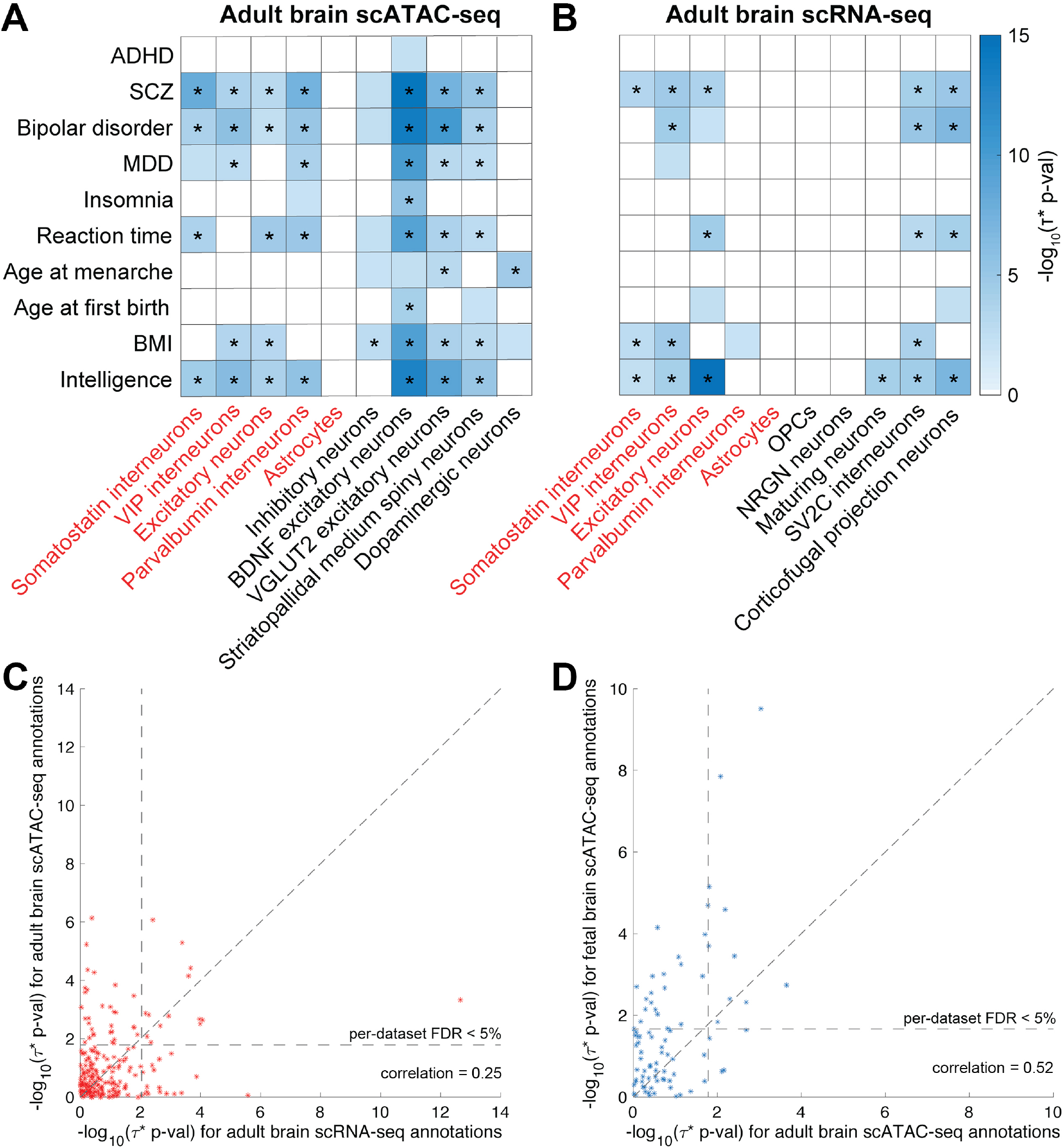
Disease enrichments of cell-type annotations derived from adult brain. We report (A) −log_10_ p-values for positive *τ* * for a subset of 10 (of 28) diseases/traits and 10 (of 18) adult brain scATAC-seq cell type annotations; (B) −log_10_ p-values for positive *τ* * for a subset of 10 (of 28) diseases/traits and 10 (of 17) adult brain scRNA-seq cell type annotations; (C) comparison of results for 8 cell types included in both adult brain scATAC-seq and scRNA-seq data; and (D) comparison of results for 3 cell types included in both fetal and adult brain scATAC-seq data. In A-B, only statistically significant results (FDR *>* 5%) are colored (−log_10_(p-value) ≥ 1.79 for scATAC-seq, ≥ 2.04 for scRNA-seq). In A-B, cell types appearing in both datasets are denoted in red font. Numerical results for all diseases/traits and cell types are reported in Table S15, Table S16, Table S17, and Table S18. * denotes Bonferroni-significant results. ADHD: attention deficit hyperactivity disorder, SCZ: schizophrenia, MDD: major depressive disorder, BMI: body mass index.

Our results also highlight disease-cell type associations that have not (to our knowledge) previously been reported in analyses of genetic data (Table 2). First, brain-derived neurotrophic factor (BDNF) excitatory neurons were highly enriched in MDD (and several other diseases/traits, including bipolar disorder and SCZ). BDNF is involved in supporting survival of existing neurons and differentiating new neurons, and decreased BDNF levels have been observed in untreated MDD^54^, bipolar ^55^ and SCZ cases ^56^. Previous studies identified an enrichment of excitatory neurons in MDD ^39^. Second, parvalbumin interneurons were enriched in bipolar disorder (and SCZ). Decreased expression and diminished function of parvalbumin interneurons in regulating balance of excitation and inhibition have been observed in bipolar disorder and SCZ cases ^57,58^. Third, vesicular glutamate transporter (VLUGT2) excitatory neurons were enriched in SCZ (as well as bipolar disorder and intelligence). VLUGT2 knock-out mice display glutamatergic deficiency, diminished maturation of pyramidal neuronal architecture, and impaired spatial learning and memory^59^, supporting a role in SCZ and intelligence.

We next analyzed adult brain scRNA-seq data spanning 17 cell types ^14^ (of which 8 were also included in the fetal brain scATAC-seq data). We identified 64 significant disease-cell type pairs (FDR *<* 5% for positive *τ* * conditional on other annotations; Table 1, Table 2, Figure 2B, Table S16). When restricting to the 33 significant disease-cell type pairs corresponding to 8 cell types included in both scATAC-seq and scRNA-seq data, 20 of 33 were also significant in analyses of scATAC-seq data. The most significant enrichment was observed for excitatory neurons in intelligence, consistent with previous genetic studies ^20^. We also identified an enrichment of corticofugal projection neurons (CPN) in intelligence, which has not (to our knowledge) previously been reported in analyses of genetic data. CPN connect neocortex and the subcortical regions and transmits axons from the cortex. Imbalance in neuronal activity, particularly regarding excitability of CPNs, has been hypothesized to lead to deficits in learning and memory ^60,61^. Recently, ref. ^62^ reported that NEUROD2 knockout mice display synaptic and physiological defects in CPN along with autism-like behavior abnormalities (where NEUROD2 is a transcription factor involved in early neuronal differentiation). CPN have previously been reported to be enriched in autism spectrum disorder (ASD) genes^63^, we did not detect a significant ASD enrichment for CPN (*p* = 0.056) or any other cell type (see Discussion).

We compared the results for 9 adult brain cell types included in both the scATAC-seq and scRNA-seq datasets (Figure 2C and Table S17). While scATAC-seq and scRNA-seq cell-type annotations for matched cell types were weakly correlated to each other (*r* = 0.01 -0.09; Table S9), the corresponding −log_10_(p-values) for positive *τ* * were moderately correlated (*r* = 0.25), confirming the shared biological information. We observed more significant p-values for scATAC-seq than for scRNA-seq in these data sets, analogous our analyses of fetal brain data (see Discussion).

We compared the results for 3 cell types (astrocytes, inhibitory neurons, excitatory neurons) included in both fetal brain and adult brain scATAC-seq data sets (Figure 2D and Table S18). While fetal brain and adult brain cell-type annotations for matched cell types were weakly correlated to each other (*r* = 0.00 - 0.01), the corresponding −log_10_(p-values) for positive *τ* * attained a moderately high correlation (*r* = 0.52), higher than the analogous correlations for scATAC-seq vs. scRNA-seq results (*r* = 0.24 for fetal brain, *r* = 0.25 for adult brain; see above). Interestingly, the enrichment in ADHD for fetal brain astrocytes (see above) was not observed for adult brain astrocytes (*p* = 0.52 for positive *τ* *, *p* = 0.0065 for difference in *τ* * for adult brain astrocytes vs. fetal brain astrocytes). While astrocytes participate in defense against stress, energy storage, and tissue repair, they also mediate synaptic pruning (elimination of synaptosomes) during development^64^. Indeed, astrocytes in more mature stages of brain development were found to be less efficient at removing synaptosomes compared to younger, fetal astrocytes ^65^ (in both in vitro in pluripotent stem cells and *in vivo* mice), supporting a fetal brain-specific role of astrocytes in brain-related diseases and traits. We also determined that the enrichment in ADHD for fetal inhibitory neurons was not observed for adult brain inhibitory neurons (*p* = 0.52 for positive *τ* *, *p* = 2.4×10^−4^ for difference in *τ* * for adult brain inhibitory neurons vs. fetal brain inhibitory neurons).

We observed little correlation between fetal brain and adult brain −log_10_(p-values) for positive *τ* * in analyses of scRNA-seq data (*r* = 0.044; Figure S2 and Table S19), possibly due to the lower power of these analyses (particularly for fetal brain scRNA-seq) in the data sets that we analyzed (see Discussion).

We performed 5 secondary analyses. First, we analyzed enrichments of both scATAC-seq and scRNA-seq brain cell types in 6 control (non-brain-related) diseases and complex traits. As expected, we did not identify any significant enrichments (Table S20 and Table S21). Second, we repeated our disease heritability enrichment analyses of scATAC-seq annotations while conditioning only on the baseline model (and not the union of open chromatin regions across all brain cell types). We identified 246 significant disease-cell type pairs, as compared to 168 significant disease-cell type pairs in our primary analysis (Figure S3A, Table S22A). This underscores the importance of conditioning on the union of open chromatin regions across all cell types, a conservative step to ensure cell-type specificity. (However, in analyses of fetal brain scATAC-seq, we obtained similar results with or without additionally conditioning on the union of open chromatin regions across all brain cell types; Figure S3B, Table S22B). Third, we performed gene set enrichment analysis using GREAT^40^ for both scATAC-seq and scRNA-seq cell-type annotations from adult brain. As expected, we identified significant enrichments in relevant gene sets (Table S23). Fourth, for the adult scRNA-seq data ^14^, we constructed annotations based on a +/-100kb window-based strategy (previously used in ref. ^8^) instead of brain-specific enhancer-gene links^7,35,36^ (used in ref. ^21^). We identified only 28 significant trait-cell type pairs (Table S24), vs. 64 using brain-specific enhancer-gene links. Fifth, we analyzed bulk chromatin data (7 chromatin marks) spanning 21 adult brain tissues^9^ (age 27-85 years). We identified 1,710 significant disease-tissue-chromatin mark triplets spanning 26 of 28 brain-related diseases and traits (Table S25). Once again, these results are included for completeness, but cannot achieve the same cell-type specificity as analyses of single-cell data.

## Discussion

We identified a rich set of disease-critical fetal and adult brain cell types by integrating GWAS summary association statistics from 28 brain-related diseases and traits with scATAC-seq and scRNA-seq data from 83 fetal and adult brain cell types ^14–17^. We confirmed many previously reported disease-cell type associations, but also identified disease-cell type associations supported by known biology that were not previously reported in analyses of genetic data. We determined that cell-type annotations derived from scATAC-seq were particularly powerful in the data that we analyzed. We also determined that the disease-cell type associations that we identified can be either shared or specific across fetal vs. adult brain developmental stages.

We note 4 key distinctions between our work and previous studies identifying disease-critical tissues and cell types ^4–8,10,12,18–21^. First, we explicitly compared results from scATAC-seq vs. scRNA-seq data in matched cell types. Although applications of single-cell data to identify disease-critical cell types have largely prioritized analyses of scRNA-seq data ^3^, we determined that cell-type annotations derived from scATAC-seq were even more powerful in our analyses. This finding may be specific to the data that we analyzed, and should not preclude further prioritization of scRNA-seq data, but does motivate further prioritization of scATAC-seq data. Second, we explicitly compared results for fetal and adult brain in matched cell types. We determined that disease-critical cell types are specific to fetal vs. adult brain developmental stages in some instances, such as the enrichment of fetal astrocytes in ADHD. Third, we rigorously conditioned on a broad set of other functional annotations, a conservative step to ensure cell-type specificity that was included in only a subset of previous studies ^39,41^. Fourth, in analyses of scRNA-seq data, we constructed annotations using brain-specific enhancer-gene links^7,35,36^ (used in ref. ^21^), an emerging approach that is more powerful than conventional window-based strategies for linking SNPs to genes.

Our findings have implications for improving our understanding of how cell-type specificity impacts disease risk. Better understanding disease-critical cell types is crucial to characterizing disease mechanisms underlying cell type specificity and developing new therapeutics ^3^. To this end, the disease-cell type associations that we identified can help guide functional follow-up experiments (e.g. Perturb-seq ^66^, saturation mutagenesis ^67^, and CRISPR-Cas9 cytosine base editor screen ^68^) to study cellular mechanisms of specific loci or genes underlying disease. In addition, our results highlight the benefits of analyzing data from different sequencing platforms and different developmental stages to identify disease-critical cell types. This motivates the prioritization of technologies that simultaneously profile ATAC and RNA expression such as SHARE-seq^69^, as well as continuing efforts to profile the developing human brain^39^.

We note several limitations of our work. First, although annotations derived from scATAC-seq generally outperformed annotations derived from scRNA-seq in the data that we analyzed, we caution that we are unable to draw any universal conclusions about which technology is most useful, as our findings may be impacted by the particularities of the data sets that we analyzed. However, we note that for both fetal and adult brain, the scRNA-seq data that we analyzed had larger numbers of donors and nuclei sequenced vs. the scATAC-seq data. Second, our resolution in identifying disease-critical cell types is fundamentally limited by the resolution of the single-cell data that we analyzed; in particular, rare but biologically important cell types may be poorly represented in these data sets. Third, despite our rigorous efforts to condition on a broad set of functional annotations, we are unable to conclude that the disease-critical cell types that we identify are biologically causal; it may often be the case that they “tag” a biologically causal cell type that is not included in the data that we analyzed. This motivates further research on methods for discriminating closely related cell types ^18^ and fine-mapping causal cell types (analogous to research on fine-mapping disease variants ^70^ and disease genes ^71^). Fourth, we failed to identify any significant cell types for 4 diseases/traits (autism, anorexia, ischemic stroke, and Alzheimer’s disease), possibly due to limited GWAS power and/or disease heterogeneity. Despite these limitations, the disease-cell type associations that we identified have high potential to improve our understanding of the biological mechanisms of complex disease.

## Data and Code Availability

The cell-type annotations and source code for primary analyses are available at https://alkesgroup.broadinstitute.org/LDSCORE/KimATAC/.

## Web Resources

S-LDSC software: https://github.com/bulik/ldsc

GWAS summary statistics: https://alkesgroup.broadinstitute.org/sumstatsformatted/

Generated cell-type annotations: https://alkesgroup.broadinstitute.org/LDSCORE/KimATAC/

Domcke et al. fetal scATAC-seq data: https://atlas.brotmanbaty.org/bbi/human-chromatin-during-development/

Cao et al. fetal scRNA-seq data: https://atlas.brotmanbaty.org/bbi/human-gene-expression-during-development/

Corces et al. scATAC-seq data: http://epigenomegateway.wustl.edu/legacy/?genome=hg38&session=drS3o1n4kJ

Velmeshev et al. scRNA-seq data: https://autism.cells.ucsc.edu/

baseline (v.1.2) annotations: https://data.broadinstitute.org/alkesgroup/LDSCORE/

1000 Genomes Project Phase 3 data: ftp://ftp.1000genomes.ebi.ac.uk/vol1/ftp/release/20130502

GREAT (Genomic Regions Enrichment of Annotations Tool): http://great.stanford.edu/

## Acknowledgements

We are grateful to Tiffany Amariuta, Katie Siewert, Martin Zhang, and Huwenbo Shi for helpful discussions. This research was funded by NIH grants U01 HG009379, U01 MH119509, R01 MH101244, R37 MH107649, R01 MH115676 and R01 MH109978. S.S.K. was supported by the NIH NHGRI award F31HG010818. This research was conducted using the UK Biobank Resource under Application 16549.

## Author Contributions

S.S.K. and A.L.P. designed experiments. S.S.K. performed experiments. K.J. and K.K.D. processed scRNA-seq data. A.Z.S assisted processing scATAC-seq data. S.R., M.K., and A.L.P. provided guidance and feedback on analyses. S.S.K. and A.L.P. wrote the manuscript with the assistance from all authors.

## Supplementary tables

See Excel file for all supplementary tables. Titles and captions are provided below.

**Table S1. List of 28 brain-related traits and 6 non-brain-related traits analyzed**. For each trait, we report a trait identifier, trait description, reference, sample size, and heritability z-score.

**Table S2. Genetic correlation among 28 brain-related traits**. We report the genetic correlation as estimated by S-LDSC across 28 GWAS summary association statistics.

**Table S3. Description of the baseline model**. For each of 53 functional annotations in the baseline model, we provide the description, proportion of SNPs annotated, and reference.

**Table S4. Tissues and cell types analyzed**. We summarize 83 cell types (for 4 single-cell datasets) and 26 tissues (for bulk chromatin data) analyzed.

**Table S5. Disease enrichments of cell-type annotations derived from fetal brain scATAC-seq data**. For each of 392 trait-cell type pairs (28 brain traits * 14 cell types), we report the proportion of SNPs, heritability enrichments, and *τ* *.

**Table S6. Matched cell types across datasets**. We report the matched cell types, defined as either exactly or closely matching cell types appearing in (A) fetal brain (scATAC-seq and scRNA-seq), (B) adult brain (scATAC-seq and scRNA-seq), (C) scATAC-seq (fetal and adult brain), and (D) scRNA-seq (fetal and adult brain).

**Table S7. Disease enrichments of cell-type annotations derived from fetal brain scRNA-seq data**. For each of 952 trait-cell type pairs (28 brain traits * 34 cell types), we report the proportion of SNPs, heritability enrichments, and *τ* *.

**Table S8. Comparison between fetal brain scATAC-seq and fetal brain scRNA-seq cell-type annotations**. For 13 cell types appearing in both fetal brain scATAC-seq and scRNA-seq datasets, we report correlation between two corresponding annotations and *τ* * (conditioning on the baseline model, union of open chromatin regions, brain-specific enhancer-gene links of all genes, and to each other).

**Table S9. Correlation among cell-type annotations**. We report the genome-wide Pearson correlation (*r*) across cell-type annotations from 4 single-cell datasets.

**Table S10. Disease enrichments of cell-type annotations derived from fetal brain scATAC-seq data in non-brain traits**. For 84 trait-cell type pairs (6 non-brain traits * 14 cell types), we report the proportion of SNPs, heritability enrichments, and *τ* *.

**Table S11. Disease enrichments of cell-type annotations derived from fetal brain scRNA-seq data in non-brain traits**. For 204 trait-cell type pairs (6 non-brain traits * 34 cell types), we report the proportion of SNPs, heritability enrichments, and *τ* *.

**Table S12. GREAT gene set enrichment of cell-type annotations derived from fetal brain datasets**. For each cell-type annotation (A) from scATAC-seq and (B) from scRNA-seq, we report the top enriched gene ontology (GO) terms, fold enrichment, binomial test p-value, and FDR-corrected q-value.

**Table S13. Disease enrichments of cell-type annotations derived from fetal brain scRNA-seq data (using window-based approach)**. We annotated SNPs +/-100kb around genes, instead of brain-specific enhancer-gene links, and assessed the heritability enrichments. For each of 952 trait-cell type pairs (28 brain traits * 34 cell types), we report the proportion of SNPs, heritability enrichments, and *τ* *.

**Table S14. Disease-critical tissues using bulk chromatin data of fetal epigenomes**. For each of 980 disease-tissue-chromatin mark triplets (28 brain traits * 5 fetal tissues * 7 chromatin marks), we report the proportion of SNPs, heritability enrichments, and *τ* *. We considered 833 high-quality epigenomes9 and selected 25 fetal biosamples related to brain tissues. We considered tier 1 assays (DNase-seq, H3K4me1, H3K4me3, H3K27ac, H3K36me3, H3K9me3 and H3K27me3), as most of observed data comes from tier 1 assays. Consistent with previous study ^7,8,11^, we built tissue annotations by annotating chromatin peaks using the MACS2^31^.

**Table S15. Disease enrichments of cell-type annotations derived from adult brain scATAC-seq data**. For each of 504 trait-cell type pairs (28 brain traits * 18 cell types), we report the proportion of SNPs, heritability enrichments, and *τ* *.

**Table S16. Disease enrichments of cell-type annotations derived from adult brain scRNA-seq data**. For each of 476 trait-cell type pairs (28 brain traits * 17 cell types), we report the proportion of SNPs, heritability enrichments, and *τ* *.

**Table S17. Comparison between adult brain scATAC-seq and adult brain scRNA-seq annotations**. For 6 cell types appearing in both adult brain scATAC-seq and scRNA-seq datasets, we report correlation between two corresponding annotations and *τ* * (conditioning on the baseline model, union of open chromatin regions, brain-specific enhancer-gene links of all genes, and to each other).

**Table S18. Comparison between fetal brain scATAC-seq and adult brain scATAC-seq annotations**. For 3 cell types appearing in both fetal brain scATAC-seq and adult brain scATAC-seq datasets, we report correlation between two corresponding annotations and *τ* * (conditioning on the baseline model, union of open chromatin regions, brain-specific enhancer-gene links of all genes, and to each other).

**Table S19. Comparison between fetal brain scRNA-seq and adult brain scRNA-seq annotations**. For 6 cell types appearing in both fetal brain scRNA-seq and adult brain scRNA-seq datasets, we report correlation between two corresponding annotations and *τ* * (conditioning on the baseline model, brain-specific enhancer-gene links of all genes, and to each other).

**Table S20. Disease enrichments of cell-type annotations derived from adult brain scATAC-seq data in non-brain traits**. For 108 trait-cell type pairs (6 non-brain traits * 18 cell types), we report the proportion of SNPs, heritability enrichments, and *τ* *.

**Table S21. Disease enrichments of cell-type annotations derived from adult brain scRNA-seq data in non-brain traits**. For 102 trait-cell type pairs (6 non-brain traits * 17 cell types), we report the proportion of SNPs, heritability enrichments, and *τ* *.

**Table S22. Disease enrichments of cell-type annotations derived from scATAC-seq data, conditioning on only baseline model**. We assessed heritability enrichments of (A) adult scATAC-seq and (B) fetal scATAC-seq cell-type annotations, conditioning on only baseline model (instead of conditioning on the baseline model and the union of chromatin marks across cell types). We report the proportion of SNPs, heritability enrichments, and *τ* *.

**Table S23. GREAT gene set enrichment of cell-type annotations derived from adult brain datasets**. For each cell-type annotation (A) from scATAC-seq and (B) from scRNA-seq, we report the top enriched gene ontology (GO) terms, fold enrichment, binomial test p-value, and FDR-corrected q-value.

**Table S24. Disease enrichments of cell-type annotations derived from adult brain scRNA-seq data (using window-based approach)**. We annotated SNPs +/-100kb around genes, instead of brain-specific enhancer-gene links, and assessed the heritability enrichments. For each of 476 trait-cell type pairs (28 brain traits * 17 cell types), we report the proportion of SNPs, heritability enrichments, and *τ* *.

**Table S25. Disease-critical tissues using bulk chromatin data of adult epigenomes**. For each of 4,116 disease-tissue-chromatin mark triplets (28 brain traits * 21 adult tissues * 7 chromatin marks), we report the proportion of SNPs, heritability enrichments, and *τ* *. We considered 32 adult biosamples related to brain tissues and tier 1 assays (DNase-seq, H3K4me1, H3K4me3, H3K27ac, H3K36me3, H3K9me3 and H3K27me3). Donor information can be found in ref. 9.

## Supplementary figures

**Figure S1.**
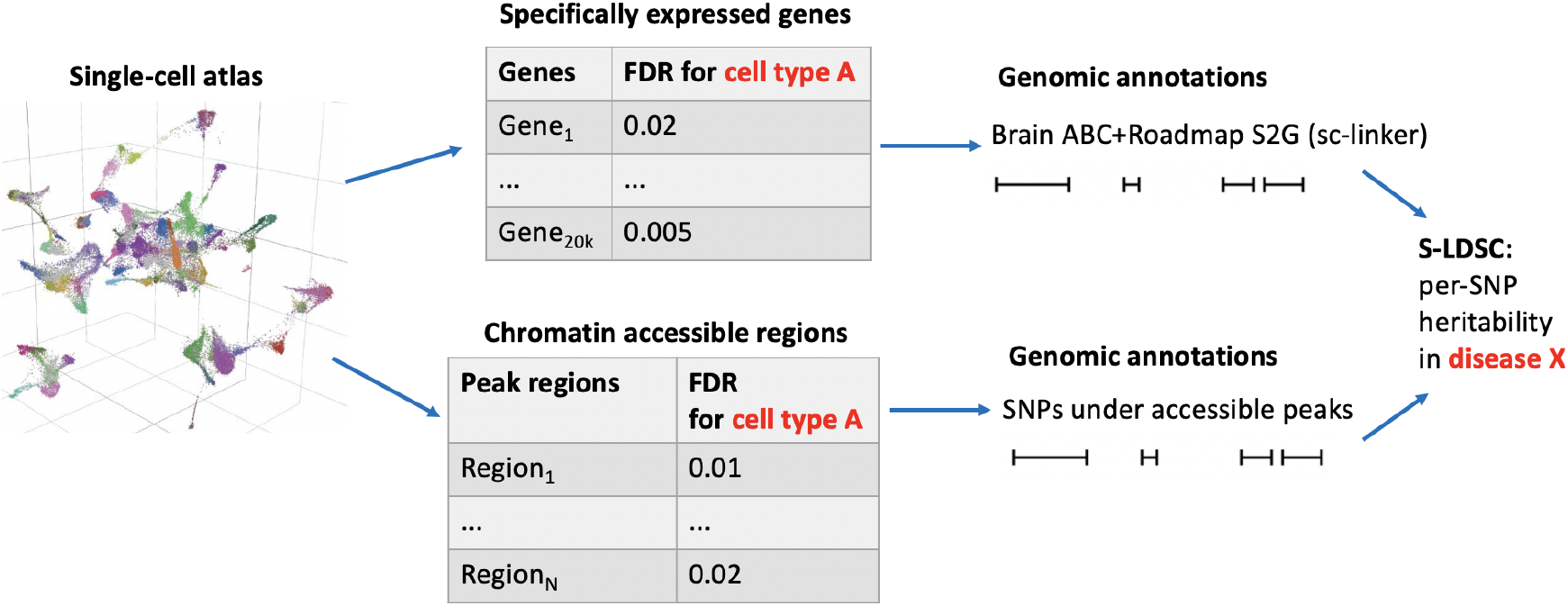
Overview of methods and analyses. We describe the overview of methods building cell-type annotations from single-cell sequencing datasets and evaluating disease informativeness applying S-LDSC across GWAS summary statistics. ABC+Roadmap S2G refers to the brain-specific SNPs-to-Genes linking strategy using enhancer-gene links ^7,21,35,36^. We separately analyzed fetal and adult brain data.

**Figure S2.**
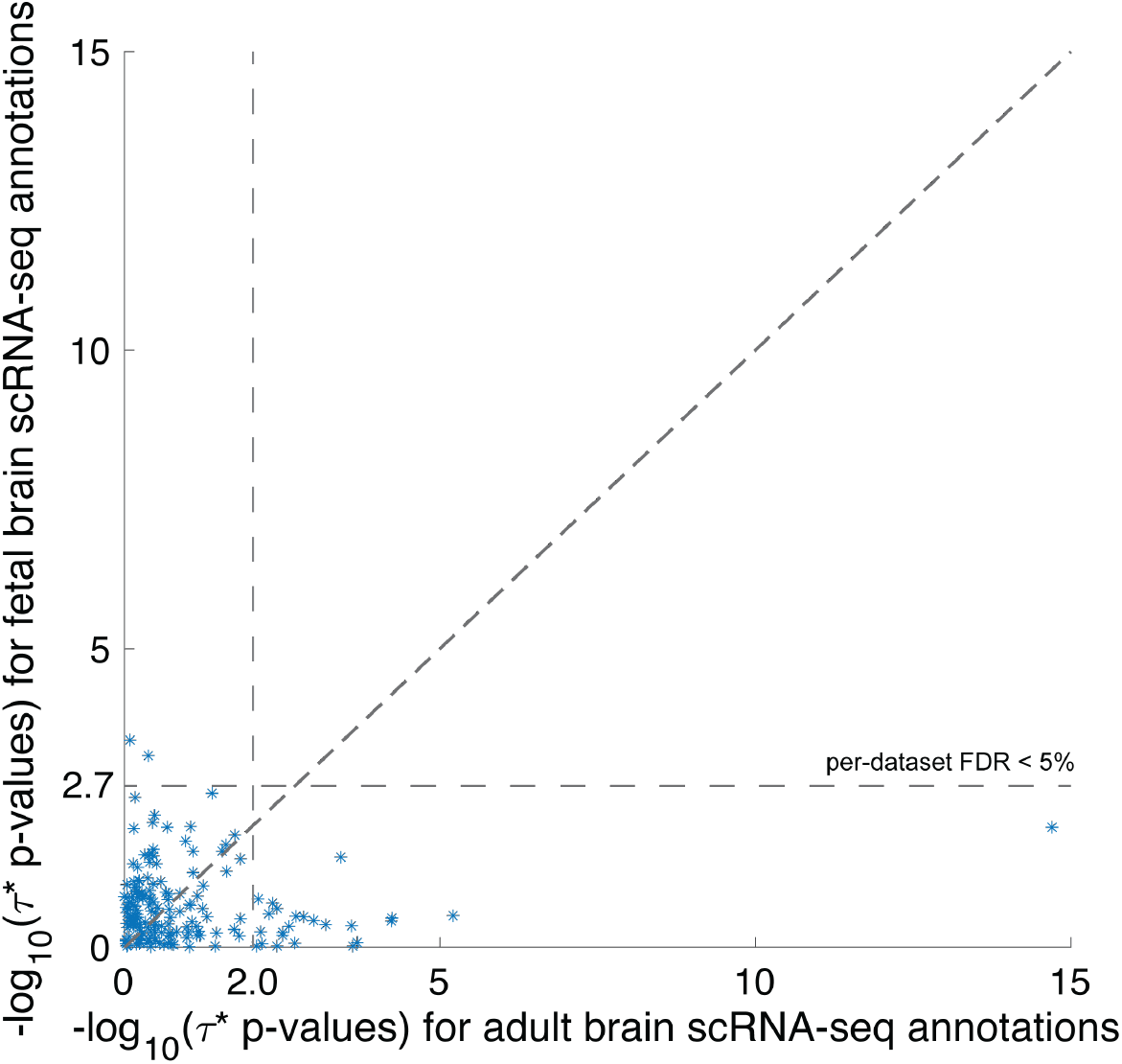
Comparison between fetal brain scRNA-seq and adult brain scRNA-seq cell-type annotations. We report −log_10_(*τ* * p-values) of fetal brain scRNA-seq and adult brain scRNA-seq annotations for 6 matched cell types (astrocytes, endothelial cells, microglia, oligodendrocytes, excitatory neurons, inhibitory neurons), conditioning on the baseline model, union of open chromatin regions, and each other. Numeric results are found in Table S19. Correlation among cell-type annotations is found in Table S9.

**Figure S3.**
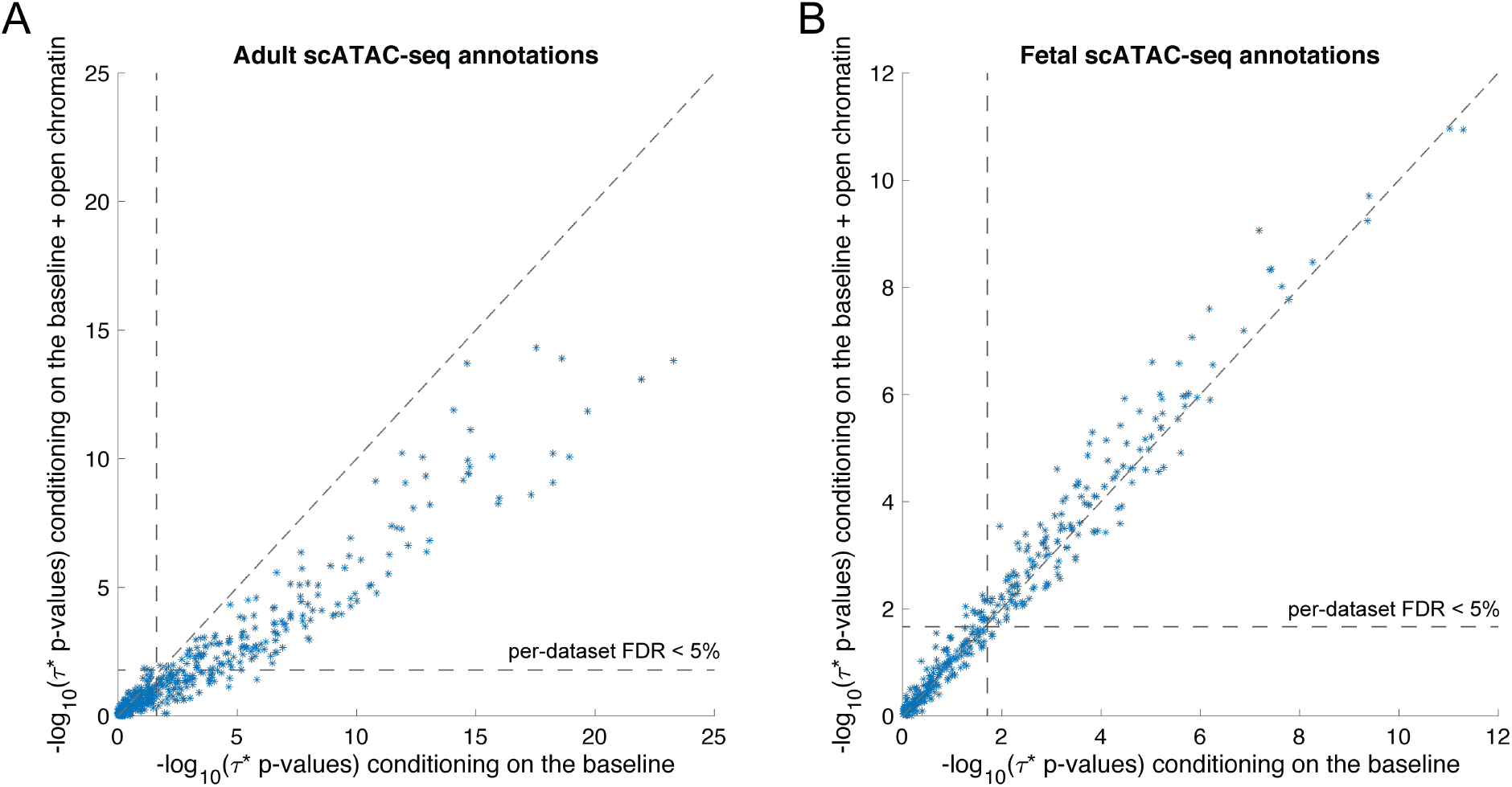
Disease enrichments of cell-type annotations derived from scATAC-seq data, conditioning on only baseline model. We report −log_10_(*τ* * p-values) of (A) adult scATAC-seq and (B) fetal scATAC-seq cell-type annotations for two different models: (1) conditioning on only baseline model and (2) baseline model and the union of chromatin marks across cell types. Numeric results are found in Table S22

## References

1. Price, A. L., Spencer, C. C., and Donnelly, P. (2015). Progress and promise in understanding the genetic basis of common diseases. Proceedings of the Royal Society B: Biological Sciences 282, 20151684.

2. Visscher, P. M., Wray, N. R., Zhang, Q., Sklar, P., McCarthy, M. I., Brown, M. A., and Yang, (2017). 10 years of gwas discovery: biology, function, and translation. The American Journal of Human Genetics 101, 5–22.

3. Hekselman, I. and Yeger-Lotem, E. (2020). Mechanisms of tissue and cell-type specificity in heritable traits and diseases. Nature Reviews Genetics 21, 137–150.

4. Maurano, M. T., Humbert, R., Rynes, E., Thurman, R. E., Haugen, E., Wang, H., Reynolds, P., Sandstrom, R., Qu, H., Brody, J., et al. (2012). Systematic localization of common disease-associated variation in regulatory dna. Science pp. 1222794.

5. Trynka, G., Sandor, C., Han, B., Xu, H., Stranger, B. E., Liu, X. S., and Raychaudhuri, S. (2013). Chromatin marks identify critical cell types for fine mapping complex trait variants. Nature Genetics 45, 124.

6. Ripke, S., Neale, B. M., Corvin, A., Walters, J. T., Farh, K.-H., Holmans, P. A., Lee, P., Bulik-Sullivan, B., Collier, D. A., Huang, H., et al. (2014). Biological insights from 108 schizophrenia-associated genetic loci. Nature 511, 421.

7. Kundaje, A., Meuleman, W., Ernst, J., Bilenky, M., Yen, A., Heravi-Moussavi, A., Kheradpour, P., Zhang, Z., Wang, J., Ziller, M. J., et al. (2015). Integrative analysis of 111 reference human epigenomes. Nature 518, 317.

8. Finucane, H. K., Reshef, Y. A., Anttila, V., Slowikowski, K., Gusev, A., Byrnes, A., Gazal, S., Loh, P.-R., Lareau, C., Shoresh, N., et al. (2018). Heritability enrichment of specifically expressed genes identifies disease-relevant tissues and cell types. Nature Genetics 50, 621.

9. Boix, C. A., James, B. T., Park, Y. P., Meuleman, W., and Kellis, M. (2021). Regulatory genomic circuitry of human disease loci by integrative epigenomics. Nature pp. 1–8.

10. Calderon, D., Bhaskar, A., Knowles, D. A., Golan, D., Raj, T., Fu, A. Q., and Pritchard, J. K. (2017). Inferring relevant cell types for complex traits by using single-cell gene expression. The American Journal of Human Genetics 101, 686–699.

11. Finucane, H. K., Bulik-Sullivan, B., Gusev, A., Trynka, G., Reshef, Y., Loh, P.-R., Anttila, V., Xu, H., Zang, C., Farh, K., et al. (2015). Partitioning heritability by functional annotation using genome-wide association summary statistics. Nature Genetics 47, 1228.

12. Ripke, S., Walters, J. T., O’Donovan, M. C., of the Psychiatric Genomics Consortium, S. W. G., et al. (2020). Mapping genomic loci prioritises genes and implicates synaptic biology in schizophrenia. MedRxiv.

13. Tanay, A. and Regev, A. (2017). Scaling single-cell genomics from phenomenology to mechanism. Nature 541, 331–338.

14. Velmeshev, D., Schirmer, L., Jung, D., Haeussler, M., Perez, Y., Mayer, S., Bhaduri, A., Goyal, N., Rowitch, D. H., and Kriegstein, A. R. (2019). Single-cell genomics identifies cell type–specific molecular changes in autism. Science 364, 685–689.

15. Cao, J., O’Day, D. R., Pliner, H. A., Kingsley, P. D., Deng, M., Daza, R. M., Zager, M. A., Aldinger, K. A., Blecher-Gonen, R., Zhang, F., et al. (2020). A human cell atlas of fetal gene expression. Science 370.

16. Domcke, S., Hill, A. J., Daza, R. M., Cao, J., O’Day, D. R., Pliner, H. A., Aldinger, K. A., Pokholok, D., Zhang, F., Milbank, J. H., et al. (2020). A human cell atlas of fetal chromatin accessibility. Science 370.

17. Corces, M. R., Shcherbina, A., Kundu, S., Gloudemans, M. J., Frésard, L., Granja, J. M., Louie, B. H., Eulalio, T., Shams, S., Bagdatli, S. T., et al. (2020). Single-cell epigenomic analyses implicate candidate causal variants at inherited risk loci for alzheimer’s and parkinson’s diseases. Nature genetics 52, 1158–1168.

18. Ulirsch, J. C., Lareau, C. A., Bao, E. L., Ludwig, L. S., Guo, M. H., Benner, C., Satpathy, T., Kartha, V. K., Salem, R. M., Hirschhorn, J. N., et al. (2019). Interrogation of human hematopoiesis at single-cell and single-variant resolution. Nature genetics 51, 683–693.

19. Hook, P. W. and McCallion, A. S. (2020). Leveraging mouse chromatin data for heritability enrichment informs common disease architecture and reveals cortical layer contributions to schizophrenia. Genome research 30, 528–539.

20. Bryois, J., Skene, N. G., Hansen, T. F., Kogelman, L. J., Watson, H. J., Liu, Z., Brueggeman, L., Breen, G., Bulik, C. M., Arenas, E., et al. (2020). Genetic identification of cell types underlying brain complex traits yields insights into the etiology of parkinson’s disease. Nature genetics 52, 482–493.

21. Jagadeesh, K. A., Dey, K. K., Motoro, D. T., Gazal, S., Engreitz, J. M., Xavier, R. J., Price, L., and Regev, A. (2021). Identifying disease-critical cell types and cellular processes across the human body by integration of single-cell profiles and human genetics. bioRxiv.

22. Kang, H. J., Kawasawa, Y. I., Cheng, F., Zhu, Y., Xu, X., Li, M., Sousa, A. M., Pletikos, M., Meyer, K. A., Sedmak, G., et al. (2011). Spatio-temporal transcriptome of the human brain. Nature 478, 483–489.

23. Pletikos, M., Sousa, A. M., Sedmak, G., Meyer, K. A., Zhu, Y., Cheng, F., Li, M., Kawasawa, Y. I., and Šestan, N. (2014). Temporal specification and bilaterality of human neocortical topographic gene expression. Neuron 81, 321–332.

24. Bakken, T. E., Miller, J. A., Ding, S.-L., Sunkin, S. M., Smith, K. A., Ng, L., Szafer, A., Dalley, R. A., Royall, J. J., Lemon, T., et al. (2016). A comprehensive transcriptional map of primate brain development. Nature 535, 367–375.

25. Li, M., Santpere, G., Kawasawa, Y. I., Evgrafov, O. V., Gulden, F. O., Pochareddy, S., Sunkin, S. M., Li, Z., Shin, Y., Zhu, Y., et al. (2018). Integrative functional genomic analysis of human brain development and neuropsychiatric risks. Science 362.

26. Mallard, T. T., Linnér, R. K., Okbay, A., Grotzinger, A. D., de Vlaming, R., Meddens, S. F. W., Tucker-Drob, E. M., Kendler, K. S., Keller, M. C., Koellinger, P. D., et al. (2020). Multivariate gwas of psychiatric disorders and their cardinal symptoms reveal two dimensions of cross-cutting genetic liabilities. BioRxiv pp. 603134.

27. Kim, S. S., Dai, C., Hormozdiari, F., van de Geijn, B., Gazal, S., Park, Y., O’Connor, L., Amariuta, T., Loh, P.-R., Finucane, H., et al. (2019). Genes with high network connectivity are enriched for disease heritability. The American Journal of Human Genetics 104, 896–913.

28. 1000 Genomes Project Consortium et al. (2015). A global reference for human genetic variation. Nature 526, 68.

29. Gazal, S., Finucane, H. K., Furlotte, N. A., Loh, P.-R., Palamara, P. F., Liu, X., Schoech, A., Bulik-Sullivan, B., Neale, B. M., Gusev, A., et al. (2017). Linkage disequilibrium–dependent architecture of human complex traits shows action of negative selection. Nature Genetics 49, 1421.

30. van de Geijn, B., Finucane, H., Gazal, S., Hormozdiari, F., Amariuta, T., Liu, X., Gusev, A., Loh, P.-R., Reshef, Y., Kichaev, G., et al. (2020). Annotations capturing cell type-specific tf binding explain a large fraction of disease heritability. Human molecular genetics 29, 1057–1067.

31. Feng, J., Liu, T., Qin, B., Zhang, Y., and Liu, X. S. (2012). Identifying chip-seq enrichment using macs. Nature protocols 7, 1728–1740.

32. Cusanovich, D. A., Hill, A. J., Aghamirzaie, D., Daza, R. M., Pliner, H. A., Berletch, J. B., Filippova, G. N., Huang, X., Christiansen, L., DeWitt, W. S., et al. (2018). A single-cell atlas of in vivo mammalian chromatin accessibility. Cell 174, 1309–1324.

33. Korsunsky, I., Millard, N., Fan, J., Slowikowski, K., Zhang, F., Wei, K., Baglaenko, Y., Brenner, M., Loh, P.-r., and Raychaudhuri, S. (2019). Fast, sensitive and accurate integration of single-cell data with harmony. Nature methods 16, 1289–1296.

34. Traag, V. A., Waltman, L., and Van Eck, N. J. (2019). From louvain to leiden: guaranteeing well-connected communities. Scientific reports 9, 1–12.

35. Fulco, C. P., Nasser, J., Jones, T. R., Munson, G., Bergman, D. T., Subramanian, V., Grossman, S. R., Anyoha, R., Doughty, B. R., Patwardhan, T. A., et al. (2019). Activity-by-contact model of enhancer–promoter regulation from thousands of crispr perturbations. Nature Genetics 51, 1664–1669.

36. Nasser, J., Bergman, D. T., Fulco, C. P., Guckelberger, P., Doughty, B. R., Patwardhan, T. A., Jones, T. R., Nguyen, T. H., Ulirsch, J. C., Lekschas, F., et al. (2021). Genome-wide enhancer maps link risk variants to disease genes. Nature pp. 1–6.

37. Hormozdiari, F., Gazal, S., van de Geijn, B., Finucane, H. K., Ju, C. J.-T., Loh, P.-R., Schoech, A., Reshef, Y., Liu, X., O’Connor, L., et al. (2018). Leveraging molecular quantitative trait loci to understand the genetic architecture of diseases and complex traits. Nature Genetics 50, 1041–1047.

38. International HapMap 3 Consortium et al. (2010). Integrating common and rare genetic variation in diverse human populations. Nature 467, 52.

39. Ziffra, R. S., Kim, C. N., Wilfert, A., Turner, T. N., Haeussler, M., Casella, A. M., Przytycki, P. F., Kreimer, A., Pollard, K. S., Ament, S. A., et al. (2020). Single cell epigenomic atlas of the developing human brain and organoids. bioRxiv pp. 2019–12.

40. McLean, C. Y., Bristor, D., Hiller, M., Clarke, S. L., Schaar, B. T., Lowe, C. B., Wenger, M., and Bejerano, G. (2010). Great improves functional interpretation of cis-regulatory regions. Nature biotechnology 28, 495–501.

41. Freimer, J. W., Shaked, O., Naqvi, S., Sinnott-Armstrong, N., Kathiria, A., Chen, A. F., Cortez, J., Greenleaf, W. J., Pritchard, J. K., and Marson, A. (2021). Systematic discovery and perturbation of regulatory genes in human t cells reveals the architecture of immune networks. bioRxiv.

42. Gazal, S., Marquez-Luna, C., Finucane, H. K., and Price, A. L. (2019). Reconciling s-ldsc and ldak functional enrichment estimates. Nature Genetics 51, 1202–1204.

43. Moriguchi, S., Takamiya, A., Noda, Y., Horita, N., Wada, M., Tsugawa, S., Plitman, E., Sano, Y., Tarumi, R., ElSalhy, M., et al. (2019). Glutamatergic neurometabolite levels in major depressive disorder: a systematic review and meta-analysis of proton magnetic resonance spectroscopy studies. Molecular psychiatry 24, 952–964.

44. Paul, K. N., Saafir, T. B., and Tosini, G. (2009). The role of retinal photoreceptors in the regulation of circadian rhythms. Reviews in endocrine and metabolic disorders 10, 271–278.

45. Sabel, B. A., Wang, J., Cárdenas-Morales, L., Faiq, M., and Heim, C. (2018). Mental stress as consequence and cause of vision loss: the dawn of psychosomatic ophthalmology for preventive and personalized medicine. EPMA journal 9, 133–160.

46. Benes, F. M. and Berretta, S. (2001). Gabaergic interneurons: implications for understanding schizophrenia and bipolar disorder. Neuropsychopharmacology 25, 1–27.

47. Dogan, B., Kazim Erol, M., Dogan, U., Habibi, M., Bulbuller, N., Turgut Coban, D., and Bulut, M. (2016). The retinal nerve fiber layer, choroidal thickness, and central macular thickness in morbid obesity: an evaluation using spectral-domain optical coherence tomography. Eur Rev Med Pharmacol Sci 20, 886–891.

48. Canto, C. B., Onuki, Y., Bruinsma, B., van der Werf, Y. D., and De Zeeuw, C. I. (2017). The sleeping cerebellum. Trends in Neurosciences 40, 309–323.

49. Batiuk, M. Y., Martirosyan, A., Wahis, J., de Vin, F., Marneffe, C., Kusserow, C., Koeppen, J., Viana, J. F., Oliveira, J. F., Voet, T., et al. (2020). Identification of region-specific astrocyte subtypes at single cell resolution. Nature communications 11, 1–15.

50. Nagai, J., Rajbhandari, A. K., Gangwani, M. R., Hachisuka, A., Coppola, G., Masmanidis, S. C., Fanselow, M. S., and Khakh, B. S. (2019). Hyperactivity with disrupted attention by activation of an astrocyte synaptogenic cue. Cell 177, 1280–1292.

51. Davies, G., Lam, M., Harris, S. E., Trampush, J. W., Luciano, M., Hill, W. D., Hagenaars, S. P., Ritchie, S. J., Marioni, R. E., Fawns-Ritchie, C., et al. (2018). Study of 300,486 individuals identifies 148 independent genetic loci influencing general cognitive function. Nature communications 9, 1–16.

52. Nirenberg, S. and Meister, M. (1997). The light response of retinal ganglion cells is truncated by a displaced amacrine circuit. Neuron 18, 637–650.

53. Li, M., Santpere, G., Kawasawa, Y. I., Evgrafov, O. V., Gulden, F. O., Pochareddy, S., Sunkin, S. M., Li, Z., Shin, Y., Zhu, Y., et al. (2018). Integrative functional genomic analysis of human brain development and neuropsychiatric risks. Science 362.

54. Lee, B.-H. and Kim, Y.-K. (2010). The roles of bdnf in the pathophysiology of major depression and in antidepressant treatment. Psychiatry investigation 7, 231.

55. Grande, I., Fries, G. R., Kunz, M., and Kapczinski, F. (2010). The role of bdnf as a mediator of neuroplasticity in bipolar disorder. Psychiatry investigation 7, 243.

56. Favalli, G., Li, J., Belmonte-de Abreu, P., Wong, A. H., and Daskalakis, Z. J. (2012). The role of bdnf in the pathophysiology and treatment of schizophrenia. Journal of psychiatric research 46, 1–11.

57. Toker, L., Mancarci, B. O., Tripathy, S., and Pavlidis, P. (2018). Transcriptomic evidence for alterations in astrocytes and parvalbumin interneurons in subjects with bipolar disorder and schizophrenia. Biological psychiatry 84, 787–796.

58. Ferguson, B. R. and Gao, W.-J. (2018). Pv interneurons: critical regulators of e/i balance for prefrontal cortex-dependent behavior and psychiatric disorders. Frontiers in neural circuits 12, 37.

59. He, H., Mahnke, A. H., Doyle, S., Fan, N., Wang, C.-C., Hall, B. J., Tang, Y.-P., Inglis, F. M., Chen, C., and Erickson, J. D. (2012). Neurodevelopmental role for vglut2 in pyramidal neuron plasticity, dendritic refinement, and in spatial learning. Journal of Neuroscience 32, 15886–15901.

60. Fernandez, F. and Garner, C. C. (2007). Over-inhibition: a model for developmental intellectual disability. Trends in neurosciences 30, 497–503.

61. Zoghbi, H. Y. and Bear, M. F. (2012). Synaptic dysfunction in neurodevelopmental disorders associated with autism and intellectual disabilities. Cold Spring Harbor perspectives in biology 4, a009886.

62. Runge, K., Mathieu, R., Bugeon, S., Lafi, S., Beurrier, C., Sahu, S., Schaller, F., Loubat, A., Herault, L., Gaillard, S., et al. (2020). Disruption of the transcription factor neurod2 causes an autism syndrome via cell-autonomous defects in cortical projection neurons. BioRxiv pp. 296889.

63. Ruzzo, E. K., Pérez-Cano, L., Jung, J.-Y., Wang, L.-k., Kashef-Haghighi, D., Hartl, C., Singh, C., Xu, J., Hoekstra, J. N., Leventhal, O., et al. (2019). Inherited and de novo genetic risk for autism impacts shared networks. Cell 178, 850–866.

64. Chung, W.-S., Clarke, L. E., Wang, G. X., Stafford, B. K., Sher, A., Chakraborty, C., Joung, J., Foo, L. C., Thompson, A., Chen, C., et al. (2013). Astrocytes mediate synapse elimination through megf10 and mertk pathways. Nature 504, 394–400.

65. Sloan, S. A., Darmanis, S., Huber, N., Khan, T. A., Birey, F., Caneda, C., Reimer, R., Quake, S. R., Barres, B. A., and Paşca, S. P. (2017). Human astrocyte maturation captured in 3d cerebral cortical spheroids derived from pluripotent stem cells. Neuron 95, 779–790.

66. Dixit, A., Parnas, O., Li, B., Chen, J., Fulco, C. P., Jerby-Arnon, L., Marjanovic, N. D., Dionne, D., Burks, T., Raychowdhury, R., et al. (2016). Perturb-seq: dissecting molecular circuits with scalable single-cell rna profiling of pooled genetic screens. Cell 167, 1853–1866.

67. Kircher, M., Xiong, C., Martin, B., Schubach, M., Inoue, F., Bell, R. J., Costello, J. F., Shendure, J., and Ahituv, N. (2019). Saturation mutagenesis of twenty disease-associated regulatory elements at single base-pair resolution. Nature communications 10, 1–15.

68. Hanna, R. E., Hegde, M., Fagre, C. R., DeWeirdt, P. C., Sangree, A. K., Szegletes, Z., Griffith, A., Feeley, M. N., Sanson, K. R., Baidi, Y., et al. (2021). Massively parallel assessment of human variants with base editor screens. Cell 184, 1064–1080.

69. Ma, S., Zhang, B., LaFave, L. M., Earl, A. S., Chiang, Z., Hu, Y., Ding, J., Brack, A., Kartha, V. K., Tay, T., et al. (2020). Chromatin potential identified by shared single-cell profiling of rna and chromatin. Cell 183, 1103–1116.

70. Schaid, D. J., Chen, W., and Larson, N. B. (2018). From genome-wide associations to candidate causal variants by statistical fine-mapping. Nature Reviews Genetics 19, 491–504.

71. Mancuso, N., Freund, M. K., Johnson, R., Shi, H., Kichaev, G., Gusev, A., and Pasaniuc, B. (2019). Probabilistic fine-mapping of transcriptome-wide association studies. Nature genetics 51, 675–682.

72. Jansen, P. R., Watanabe, K., Stringer, S., Skene, N., Bryois, J., Hammerschlag, A. R., de Leeuw, C. A., Benjamins, J. S., Muñoz-Manchado, A. B., Nagel, M., et al. (2019). Genome-wide analysis of insomnia in 1,331,010 individuals identifies new risk loci and functional pathways. Nature genetics 51, 394–403.

73. Loh, P.-R., Kichaev, G., Gazal, S., Schoech, A. P., and Price, A. L. (2018). Mixed-model association for biobank-scale datasets. Nature Genetics 50, 906–908.

74. Pardiñas, A. F., Holmans, P., Pocklington, A. J., Escott-Price, V., Ripke, S., Carrera, N., Legge, S. E., Bishop, S., Cameron, D., Hamshere, M. L., et al. (2018). Common schizophrenia alleles are enriched in mutation-intolerant genes and in regions under strong background selection. Nature Genetics 50, 381.

75. Dashti, H. S., Jones, S. E., Wood, A. R., Lane, J. M., Van Hees, V. T., Wang, H., Rhodes, J. A., Song, Y., Patel, K., Anderson, S. G., et al. (2019). Genome-wide association study identifies genetic loci for self-reported habitual sleep duration supported by accelerometer-derived estimates. Nature communications 10, 1–12.

76. Demontis, D., Walters, R. K., Martin, J., Mattheisen, M., Als, T. D., Agerbo, E., Baldursson, G., Belliveau, R., Bybjerg-Grauholm, J., Bækvad-Hansen, M., et al. (2019). Discovery of the first genome-wide significant risk loci for attention deficit/hyperactivity disorder. Nature genetics 51, 63–75.

77. Howard, D. M., Adams, M. J., Clarke, T.-K., Hafferty, J. D., Gibson, J., Shirali, M., Coleman, J. R., Hagenaars, S. P., Ward, J., Wigmore, E. M., et al. (2019). Genome-wide meta-analysis of depression identifies 102 independent variants and highlights the importance of the prefrontal brain regions. Nature neuroscience 22, 343–352.

78. Stahl, E. A., Breen, G., Forstner, A. J., McQuillin, A., Ripke, S., Trubetskoy, V., Mattheisen, M., Wang, Y., Coleman, J. R., Gaspar, H. A., et al. (2019). Genome-wide association study identifies 30 loci associated with bipolar disorder. Nature genetics 51, 793–803.

79. Savage, J. E., Jansen, P. R., Stringer, S., Watanabe, K., Bryois, J., De Leeuw, C. A., Nagel, M., Awasthi, S., Barr, P. B., Coleman, J. R., et al. (2018). Genome-wide association meta-analysis in 269,867 individuals identifies new genetic and functional links to intelligence. Nature genetics 50, 912–919.

